# *P. aeruginosa* liquid-based pathogenesis triggers HLH-30-dependent metabolic rewiring in *C. elegans*

**DOI:** 10.64898/2026.09.03.749215

**Authors:** Lois A. Armendariz, Elissa Tjahjono, Yvette Acevedo, Fernanda Vaca, Jane Lowder, Anna Singh, Nikita Singh, Alexey V. Revtovich, Ainsley High, Evan Park, Natalia V. Kirienko

**Affiliations:** Department of BioSciences, 6100 Main St, MS140, Rice University, Houston TX 77005 USA

## Abstract

Innate immunity is the first line of defense against invading pathogens and is essential for maintaining host survival. While the majority of innate immunity studies have focused on pathogen recognition and antimicrobial responses, increasing evidence suggests that lipid metabolism plays a fundamental role in shaping immune function. Understanding how these metabolic pathways contribute to immunity is crucial in the context of bacterial infections caused by opportunistic pathogens such as *Pseudomonas aeruginosa*. Host defense against *P. aeruginosa* requires the coordination of innate immune and metabolic responses; however, the mechanisms linking lipid metabolism to pathogen resistance remain poorly understood. Research into the relationship between lipid homeostasis and innate immunity may reveal factors governing host-pathogen interactions and identify novel strategies to enhance resistance to infection.

Here, we demonstrate that *P. aeruginosa* liquid-based pathogenesis (LK-*Pa*) triggers a shift in host metabolism which differs from the one observed in response to agar-based pathogenesis. Our bioinformatic analyses revealed a highly similar metabolic profile (enrichment of lipid metabolism) in worms exposed to LK-*Pa* or the iron chelator phenanthroline, suggesting a shared host response to iron deprivation. We further characterized the host genetic factors driving this metabolic shift and established their importance for host defense against LK-*Pa* as well as liquid-based pathogenesis by Gram-positive pathogens *Enterococcus faecalis* and *Staphylococcus aureus*. Notably, our results indicate that LK-*Pa* triggers host lipid droplet depletion, an upstream component that leads to increased β-oxidation. Finally, we demonstrate that LK-*Pa* triggers repression of MXL-3 which results in HLH-30-dependent metabolic rewiring.

**Author Summary:** Pathogenesis alters host metabolism in ways that can either enhance survival or promote disease. Understanding how pathogenesis triggers host metabolic rewiring may reveal novel strategies to combat antimicrobial-resistant infections. Using *Caenorhabditis elegans*, we studied how the opportunistic pathogen *P. aeruginosa* reshapes host metabolism during liquid-based infection.

Our results reveal that liquid-based pathogenesis induces extensive activation of host lipid metabolism, a distinct transcriptional response from the one triggered during agar-based pathogenesis. This lipid signature closely resembles the host response to iron deprivation, suggesting that in the case of *P. aeruginosa*, bacterial siderophore production is a major driver of host metabolic rewiring. We identified seventeen lipid genes crucial for survival during *P. aeruginosa* exposure in liquid; most of them were also needed for defense against the Gram-positive pathogens *E. faecalis* and *S. aureus*. Notably, most of these genes function independently of activating known organellar surveillance pathways.

Instead, we show that *P. aeruginosa* liquid-based pathogenesis triggers host lipid droplet depletion and induces a metabolic shift towards β-oxidation. This process is driven by inhibition of the nutrient-responsive transcription factor MXL-3 and subsequent activation of the lysosomal transcription factor HLH-30, which drives expression of β-oxidation genes. Altogether, our findings identify an MXL-3 -HLH-30 regulatory axis that couples pathogen-induced metabolic stress to lipid mobilization and reveal metabolic rewiring as a host defense mechanism during bacterial infection.

## Introduction

Living organisms are, by definition, an invaluable source of biologically useful materials. This makes them an ideal target for pathogens, which infect other organisms to access the “goodies” within. The fight for these materials has led to a complex and dynamic evolutionary arms race between hosts and pathogens. One example of this, and a current, pressing threat to human health, is the rapid increase and global spread of antimicrobial resistance (AMR)^1,2^. Misuse of antibiotics by the health and agricultural industries has accelerated the rate at which microbes acquire resistance determinants^1,3^. Solutions involving the development of new antimicrobials have been unable to slow AMR, possibly because of low pharmaceutical investments.

Multidrug-resistant (MDR) infections have been on the rise within hospitals, which are hot spots for sick and immunocompromised people^4^. In hospital settings, *Pseudomonas aeruginosa*, *Enterococcus faecalis*, and *Staphylococcus aureus* are particularly problematic^5^. *P. aeruginosa* is a Gram-negative opportunistic pathogen primarily associated with pneumonia, urinary tract infections, and bloodstream infections; it is also the main cause of morbidity and mortality in patients with cystic fibrosis^6^. *P. aeruginosa* exhibits intrinsic resistance mechanisms, biofilm formation, and a rapid acquisition of resistance determinants, all of which can result in MDR isolates^6^. *E. faecalis*, a Gram-positive commensal normally found in the gastrointestinal tract, is now a leading cause of hospital-acquired urinary tract infections, bloodstream infections, and endocarditis^7^. Similarly, *S. aureus*, a Gram-positive commensal commonly found on the skin, has become a major cause of soft tissue and skin infections, sepsis, and pneumonia^8^.

Previous research in *Caenorhabditis elegans* has shown that lipid metabolism plays a crucial role in host defense regulation^9,10^. Beyond their canonical function in energy storage and membrane biosynthesis, lipids can act as signaling molecules influencing innate immunity, stress responses, and pathogen resistance^11,12^. Studies have established that transcriptional regulators of lipid metabolism such as NHR-49, MDT-15, and HLH-30 are required for defense against agar-based (chronic) bacterial infections^13–16^. Furthermore, perturbations in fatty acid desaturation and lipid storage can significantly alter a host’s ability to survive infection, highlighting metabolic rewiring as a crucial component of the innate immune response^13–17^. Notably, infection triggers extensive host lipid metabolic changes, suggesting that organisms actively reprogram their metabolic profile to meet the demands imposed by pathogenesis^16,17^. Despite these significant advances, the mechanisms by which lipid metabolism contributes to the response to liquid-based pathogenesis (acute infection) remain poorly understood.

Lipid metabolism has also been linked to the regulation of organellar surveillance during stress and infection^9,10,13,18^. In *C. elegans*, dysregulated lipid homeostasis can affect the function of multiple organelles, such as mitochondria, endoplasmic reticulum, lysosomes, and lipid droplets^19–22^. These organelles are crucial for mounting cellular stress responses and innate immunity. Our lab has highlighted the role of lipid metabolism in the regulation of mitochondrial surveillance pathways such as the mitochondrial unfolded protein response (UPR^mt^) and the Ethanol and Stress Response Element (ESRE) network^10,17^. Lipid droplets have been shown to contribute to host defense by serving as reservoirs of metabolic substrates and immune effectors, while lysosomal signaling and autophagy pathways are tightly coupled to lipid availability and mobilization^23,24^. Altogether, these studies suggest that lipid metabolism is at the center of organellar homeostasis and immune defense, coordinating adaptive responses to pathogenic stress. Understanding how lipid metabolic pathways interact with organellar surveillance pathways may reveal fundamental principles governing the host response to infection and help identify new targets for combating AMR pathogens.

In this study, we reveal that liquid-based *P. aeruginosa* pathogenesis (LK-*Pa*) triggers a host response characterized by extensive activation of lipid metabolism and β-oxidation, which is in part caused by with siderophore-mediated iron deprivation. We identified seventeen lipid genes crucial for host survival against liquid-based exposure to Gram-negative and Gram-positive pathogens. We further show that LK-*Pa* triggers host lipid droplet depletion and a metabolic shift towards β-oxidation. Finally, we reveal an MXL-3–HLH-30 regulatory pathway that links pathogen-induced metabolic stress to lipid mobilization and β-oxidation, highlighting metabolic rewiring as a host defense mechanism during acute infection.

## Results

### Lipid metabolism is enriched during P. aeruginosa liquid-based pathogenesis

Our lab previously established that there is a difference in host gene expression between *P. aeruginosa* liquid-based (referred to as LK-*Pa*) and agar-based (referred to as SK-*Pa*) pathogenesis^25^. To understand the host response differences driven by these two conditions, fold changes in gene expression in LK-*Pa* were normalized to those in SK-*Pa*, thereby contrasting genes that are differentially regulated (>2-fold) by exposure to the pathogen in each context. A similar comparison of gene expression in worms exposed to the common laboratory food *E. coli* (OP50) in both settings (liquid-based: LK-*Ec* and agar-based: SK-*Ec*) was performed to account for environmentally driven changes in gene expression. A principal component analysis (PCA) performed on a total of 2,783 differentially regulated genes in these four conditions revealed contextual differences in host response patterns (**Fig. 1A**).

**Figure 1.**
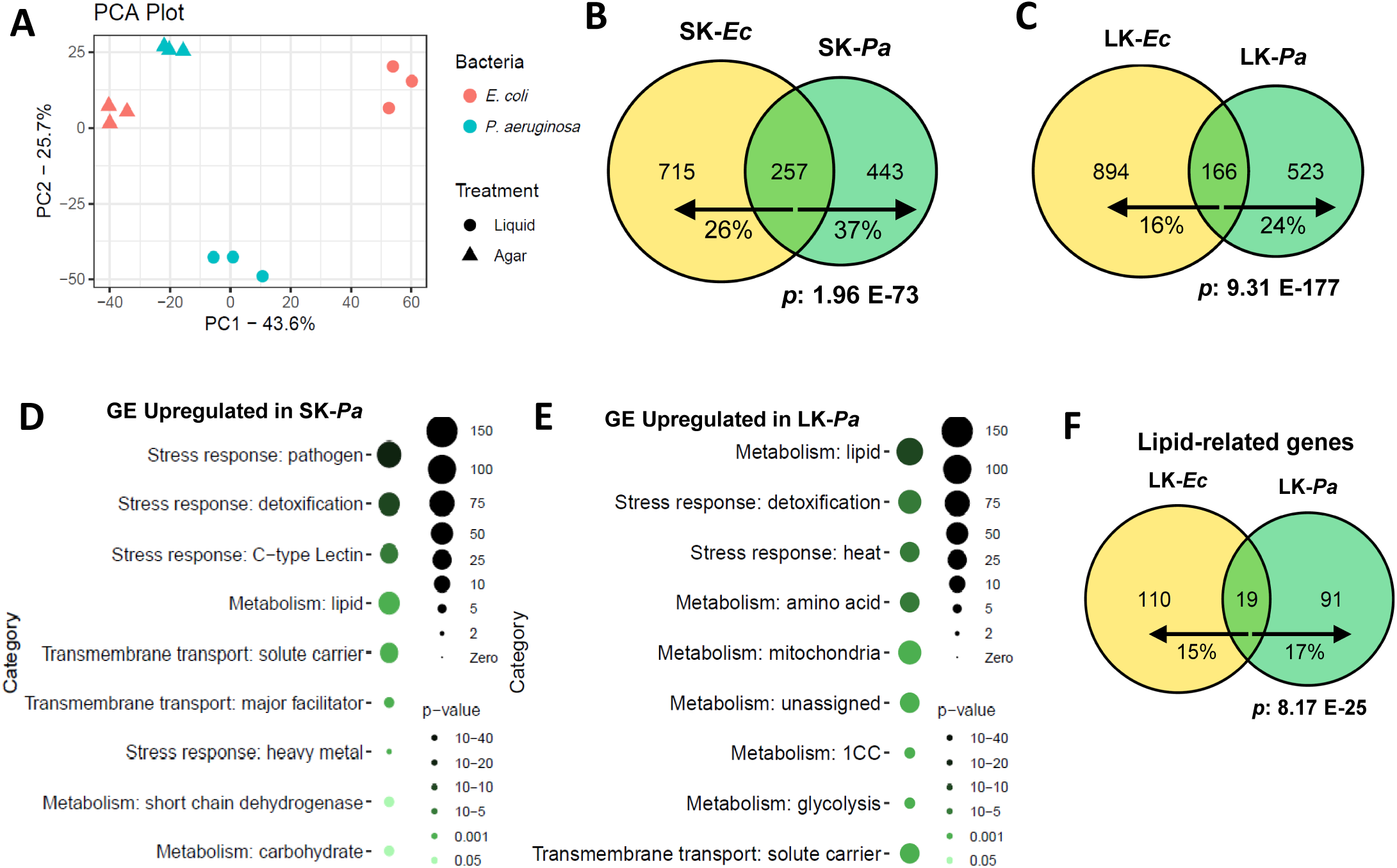
*P. aeruginosa* liquid-based pathogenesis changes host lipid metabolism. **(A)** Principal component analysis of differentially regulated genes in response to LK-*Ec*, LK-*Pa*, SK-*Ec*, and SK-*Pa*. **(B, C, F)** Venn diagrams of genes upregulated in response to SK-*Ec* and SK-*Pa,* **(B)** LK-*Ec* and LK-*Pa,* **(C)** or lipid genes upregulated in response to LK-*Ec* and LK-*Pa* **(F)**. **(D-E)** WormCat gene enrichment analysis of genes upregulated in response to SK-*Pa* **(D)** or LK-*Pa* **(E)**. In all Venn diagrams, arrows point toward the treatment for which the corresponding percentage of shared genes (shown below each arrow) was calculated.

Comparisons of gene expression between SK-*Pa* and its non-pathogenic counterpart, SK-*Ec*, showed large overlap **(Fig. 1B)**. A total of 972 and 700 genes were upregulated in response to SK-*Ec* and SK-*Pa*, respectively, while 257 genes were shared between the two (1,415 unique genes total). Comparisons between LK-*Pa* and LK-*Ec* showed a similar, if less overt, pattern **(Fig. 1C)**. Out of a total of 1,583 genes, 166 were shared, indicating a larger difference between pathogenesis and growth in liquid than on agar. Gene enrichment analyses were performed on differentially regulated genes from SK-*Pa* **(Fig. 1D)** and LK-*Pa* **(Fig. 1E)** using WormCat^26^. As expected, in response to SK-*Pa*, *C. elegans* activated effectors mapping to pathogenic and detoxification stress responses. Unexpectedly, the most enriched biological processes in response to LK-*Pa* were lipid metabolism and detoxification stress response. Gene enrichment analysis on differentially regulated genes in response to growth on *E. coli* revealed enrichment of baseline metabolism pathways **(Fig. S1A-B).**

To investigate whether the change in lipid metabolism was in response to LK-*Pa* vs. the environment, we used Wormcat^26^ and DAVID^27,28^ to identify lipid-related genes upregulated in LK-*Ec* (129 genes) and LK-*Pa* (110 genes). Only 19 were shared between the two **(Fig. 1F)**, suggesting that the majority of changes to lipid metabolism in LK-*Pa* were due to *P. aeruginosa* virulence in liquid rather than adaptation to the liquid environment.

### Iron deprivation triggers changes in lipid metabolism

Previously, we demonstrated that exposure to the iron chelator 1,10-phenanthroline closely mimics LK-*Pa*, where iron removal is mediated by the siderophore pyoverdine. When gene expression data of worms treated with phenanthroline were included in the PCA, a similar host-response pattern to that of LK-*Pa* was observed, showing a strong separation from agar conditions or LK-*Ec* in PC2 (**Fig. 2A**). A closer gene enrichment analysis towards each principal component’s top contributors was performed by collecting 300 genes with the largest loading scores for both positive and negative directions (**Table S1**). All 300 genes passed the “importance” cutoff as determined by √1/*n*, where *n* equals the number of genes. For PC1, the negative loading score contributors (particularly profound in SK-*Pa*) mainly played a role in general metabolic adaptation. Meanwhile, positive loading contributors in PC1 (predominately in LK-*Ec*) were enriched for genes that increase translation, likely driven by adaptation to the liquid environment. For PC2, negative loading score contributors separated LK-*Pa* and phenanthroline conditions from the rest and showed enrichment in lipid metabolic processes such as β-oxidation, amino acid breakdown, as well as flavin-containing monooxygenases (FMO). This fits a typical mitochondrial damage and hypoxic response that occurs due to iron deprivation^29,30^. Positive loading score contributors for PC2 were enriched for stress response genes, likely driven by the pathogenesis response in SK-*Pa*.

**Figure 2.**
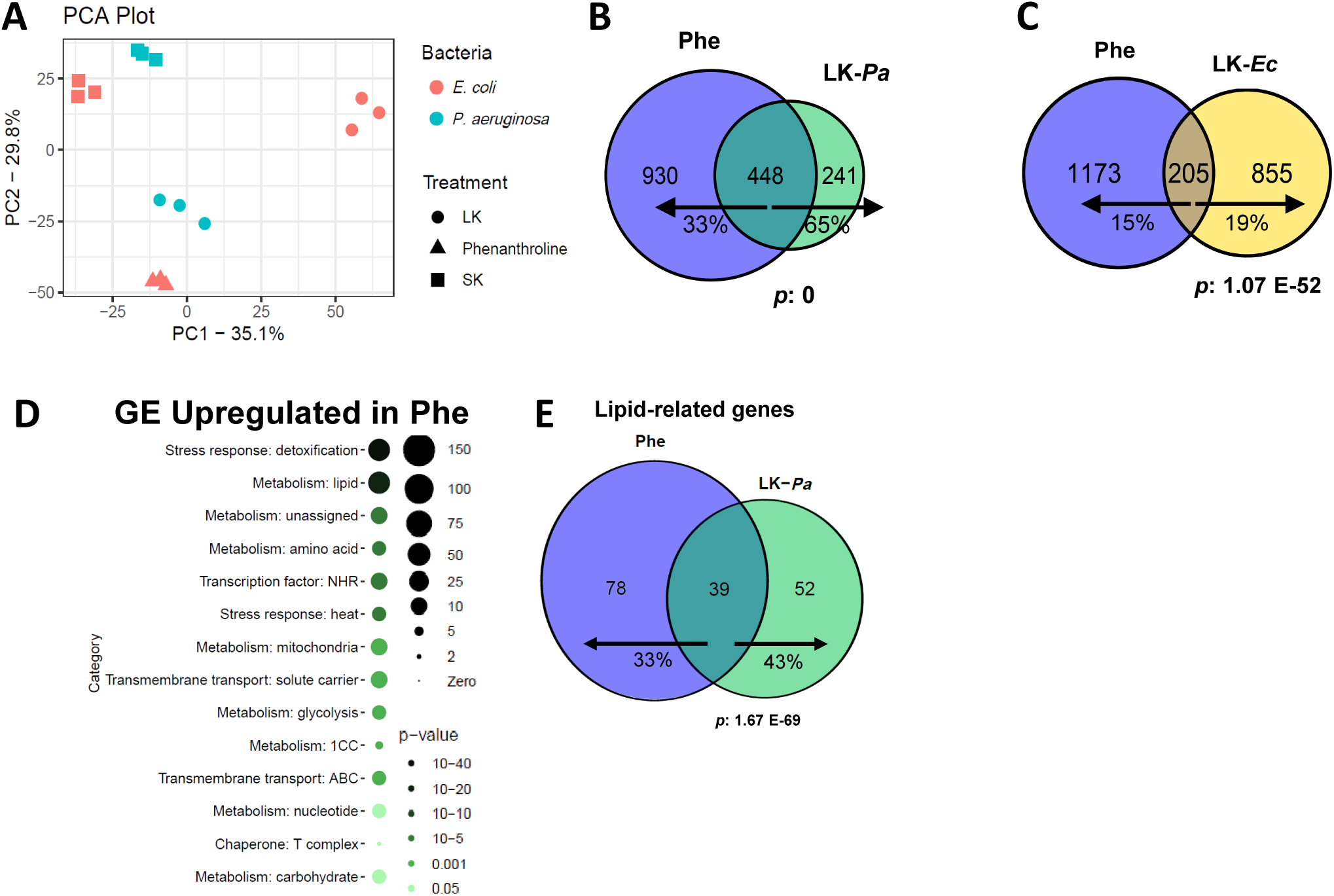
Iron starvation rewires host lipid metabolism. **(A)** Principal component analysis of differentially regulated genes in response to LK-*Ec*, LK-*Pa*, SK-*Ec*, SK-*Pa*, and phenanthroline in liquid. **(B, C, E)** Venn diagrams of genes upregulated in response to phenanthroline and LK-*Pa*, **(B)** phenanthroline and LK-*Ec*, **(C)** or lipid genes upregulated in response to phenanthroline and LK-*Pa*. **(D)** Wormcat gene enrichment analysis of genes upregulated in response to phenanthroline. In all Venn diagrams, arrows point toward the treatment for which the corresponding percentage of shared genes (shown below each arrow) was calculated.

A total of 1,378 and 689 genes were upregulated in response to phenanthroline and LK-*Pa*, respectively **(Fig. 2B)**. A large proportion of these genes (448) were shared, suggesting that iron deprivation plays a major role in these responses and drives ∼65% of transcriptional changes. A much smaller overlap of 205 genes was observed between LK-*Ec* and phenanthroline, likely indicating genes that are influenced by environmental adaptation (**Fig. 2C**). Notably, gene enrichment analysis of the phenanthroline condition showed significant enrichment for detoxification stress response and lipid metabolism, similar to LK-*Pa* **(Fig. 2D)**. A total 117 lipid-related genes were upregulated by phenanthroline, and 39 of these overlapped with the 91 lipid genes upregulated by LK-*Pa*, strengthening gene enrichment results **(Fig 2E)**.

### Lipid metabolism is required for host defense against in liquid

Further investigation into the metabolic categories driving the significance of the gene enrichment analysis revealed that LK-*Pa* upregulated a large number of β-oxidation genes. A total of 20 genes involved in β-oxidation were upregulated during LK-*Pa* exposure, as compared to 7 genes during SK-*Pa* exposure **(Fig. 3A-B)**. Because lipid metabolism was the most enriched category in the LK-*Pa* condition, we hypothesized that lipid metabolism genes may play a role in host defense. To investigate this, we conducted an RNAi screen on the 91 lipid-related genes upregulated by LK-*Pa* and evaluated whether their knockdown decreased worm survival in LK-*Pa* treatment. Out of the 86 genes that were screened (5 were not screened due to the lack of RNAi construct), eighteen were required for survival in the LK-*Pa* assay **(Fig. 3C)**. To assess whether the increase in susceptibility was due to pathogen exposure rather than an effect on worm lifespan, we screened these eighteen hits in LK-*Ec* conditions **(Fig. 3D)**. Only knockdown of *C32D5.12* resulted in increased death in LK-*Ec*. This was unsurprising, since prior research showed *C32D5.12(RNAi)* induced larval arrest^31^. Additionally, it was shown that it potentially affects locomotion, a process necessary for survival in a liquid environment^32^. As such, *C32D5.12* was deemed non-specific and was removed from the panel.

**Figure 3.**
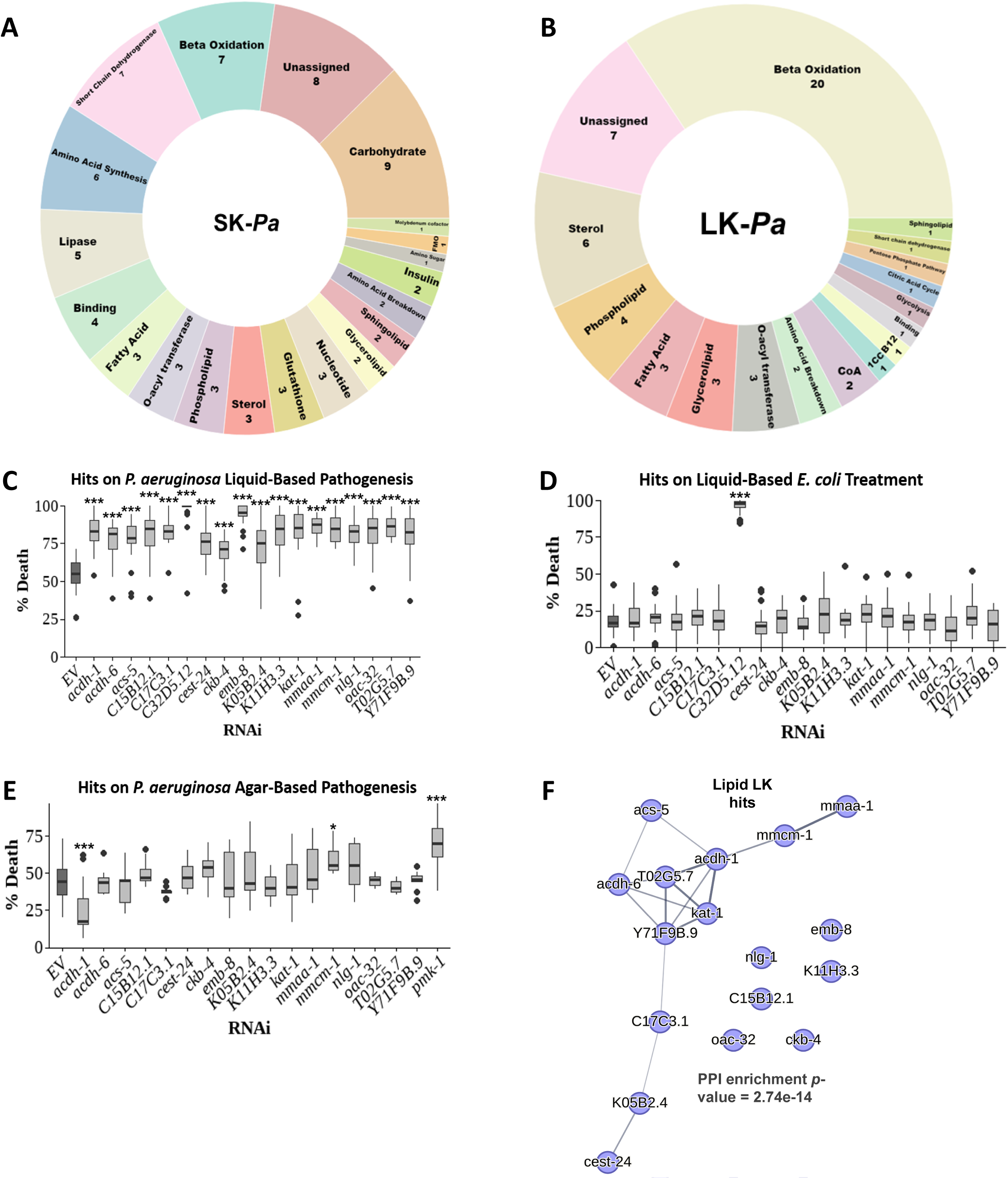
Disruption of lipid metabolism sensitizes *C. elegans* to *P. aeruginosa* in a liquid-killing model (A-B) Sunburst charts categorizing the pathways of lipid-related genes upregulated in SK-*Pa* **(A)** or LK-*Pa* **(B)**. **(C-E)** Quantification of the percentage of dead worms for different knockdowns during LK-*Pa* **(C)**, LK-*Ec* **(D)**, or SK-*Pa* **(E)**. One-way analysis of variance (ANOVA) was performed to calculate the significance of a treatment or a condition when there were more than two cohorts in the experimental setting. No stars = not significant, \**p* < 0.05, and *** *p* < 0.001. **(F)** STRING analysis for protein-protein interactions; line thickness indicates the strength of data support for the connected nodes.

Subsequently, knockdowns of the remaining seventeen hits were evaluated for their effect on worm survival in SK-*Pa* **(Fig. 3E)**. Out of these, the majority had no effect, *mmcm-1(RNAi)* increased susceptibility to *P. aeruginosa* agar-based pathogenesis, while *acdh-1(RNAi)* promoted resistance. The sensitivity of *mmcm-1(RNAi)* to SK-*Pa* could be due to increased levels of propionate, a toxic metabolite in the vitamin B12-dependent pathway^33–35^. Meanwhile, the resistance of *acdh-1(RNAi)* to SK-*Pa* could be due to activation of the canonical B12-dependent pathway, rather than activation of the shunt pathway^36^. This is consistent with our prior research showing the importance of B12 sufficiency for survival in LK-*Pa*^37^. To better understand the relationship between the seventeen hits, STRING analysis was performed to evaluate protein-protein interactions^38^ **(Fig. 3F)**. Our results reveal a prominent mitochondrial β-oxidation/metabolism module comprised of ACS-5, ACDH-1, ACDH-6, KAT-1, MMAA-1, and MMCM-1, which are strongly functionally associated. There were also some proteins forming disconnected nodes, suggesting a distinct pathway influencing host defense in contrast to the core β-oxidation network.

### Lipid metabolism is required for host defense against Gram-positive pathogens in liquid

Given that lipid metabolism genes play a role in host defense in LK-*Pa*, their effect on survival in liquid-based pathogenesis by Gram-positive pathogens was also evaluated. To investigate this, knockdowns of each of the seventeen lipid genes were exposed to *Enterococcus faecalis (Ef)* or *Staphylococcus aureus (Sa)* in liquid. Worm death was normalized across each pathogenesis model by computing *ΔDeath*, which represents the difference in mean percent death between each RNAi knockdown and the *EV(RNAi)* control within each pathogenesis model. Thus, a positive *ΔDeath* value would indicate higher susceptibility relative to control worms, and near-zero values indicate little to no susceptibility towards a pathogen.

To assess whether susceptibility phenotypes were similar across pathogenesis models, pairwise Spearman correlations were computed on the *ΔDeath* for all RNAi knockdowns across. There was a strong positive correlation between the Gram-positive pathogens *E. faecalis* and *S. aureus* (r_s_ = 0.86, *p* = 4.8e^-06^), but no significant correlation was observed between Gram-positive and Gram-negative pathogens (r_s_ = 0.26, *p* = 0.30 *Pa* to *Ef* and r_s_ = 0.28, *p* = 0.27 *Pa* to *Sa*). The triangle map shows the relative contribution of each pathogen to susceptibility across knockdowns **(Fig. 4A)**. Each vertex represents maximum susceptibility towards a pathogen. Therefore, knockdowns located near a vertex demonstrate high susceptibility to that specific pathogen, whereas location near an edge indicates shared susceptibility between the two adjacent pathogens. Knockdowns located at the center of the triangle exhibit broad susceptibility across all three pathogens. The size of a circle indicates the magnitude, which represents the effect on host survival across all three pathogens for each knockdown (a larger circle indicates a more prominent effect on host survival across all conditions). Lastly, the color assigned to each knockdown represents the pathogen it is most susceptible to. A stacked susceptibility bar graph complements the triangle map by presenting the absolute *ΔDeath* induced by each pathogen for a specific knockdown **(Fig. 4B)**. All the knockdowns tested had an increase in death on Gram-positive bacteria compared to the *EV(RNAi)* control.

**Figure 4.**
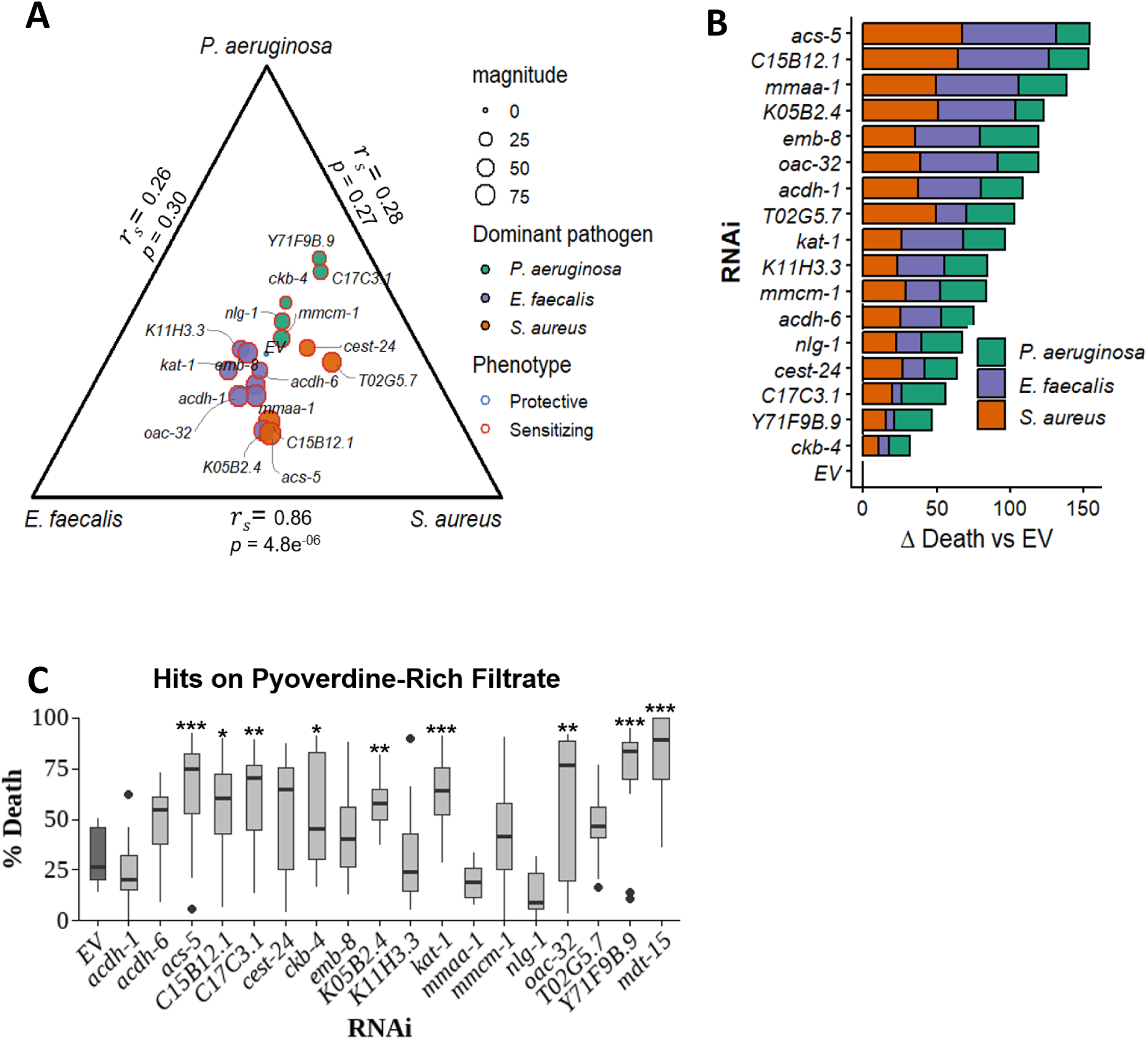
Knockdown of lipid metabolic genes increases susceptibility to liquid-based *E. faecalis* and *S. aureus* pathogenesis. **(A)**Ternary plot (triangle map) showing the relative contribution of RNAi-mediated susceptibility phenotypes across liquid pathogenesis by *P. aeruginosa*, *E. faecalis*, and *S. aureus*. Circle size for each RNAi corresponds to susceptibility magnitude, calculated as the sum of the absolute *ΔDeath* values across all pathogens. Spearman correlation coefficients are located on the edges, representing pairwise comparisons between the two neighboring pathogens. **(B)** Stacked bar plot representing the susceptibility of each RNAi across all pathogens. Bars represent *ΔDeath* relative to *EV(RNAi)*. Total bar length reflects the overall magnitude of the susceptibility phenotype, while colored segments indicate the contribution of each pathogen to the total phenotype. **(C)** Quantification of the percentage of dead worms for different knockdowns during treatment with pyoverdine-rich filtrate derived from *P. aeruginosa*. One-way analysis of variance (ANOVA) was performed to calculate the significance of a treatment or a condition when there were more than two cohorts in the experimental setting. No stars = not significant, * *p* < 0.05, ** *p* < 0.01, and *** *p* < 0.001.

We hypothesized that the production of pyoverdine by *P. aeruginosa* could be a main driver for the lack of correlation between Gram-negative and Gram-positive pathogens. To investigate this, knockdowns of the seventeen lipid gene hits were exposed to *P. aeruginosa*-free spent media containing large amounts of pyoverdine (hereafter referred to as filtrate), and the resulting death was measured **(Fig. 4C)**. More than half of the knockdowns showed higher death than the *EV(RNAi)* control, indicating that pyoverdine plays a major role in triggering these changes in host lipid metabolism.

### Lipid metabolism impact on host defense is independent of organellar surveillance pathways

Previous research in our lab has identified the mitochondrial ethanol and stress response element (ESRE) as a host response mechanism to LK-*Pa*^25^. We showed that when *C. elegans* is exposed to *P. aeruginosa* in a liquid milieu, the pathogen scavenges iron by secreting pyoverdine. This siderophore damages host mitochondria and triggers activation of ESRE and selective autophagic degradation of mitochondria through the PINK-1/Parkin pathway if mitochondrial damage is unresolved^25,30,39^. Others have shown that LK-*Pa* can trigger activation of UPR^mt^ effectors^40^. Altogether, these studies highlight the importance of organellar surveillance and proteostasis in response to liquid-based pathogenesis by *P. aeruginosa*. For this reason, we investigated the role of our lipid metabolism gene hits in the activation of organellar surveillance.

To study their effect on ESRE activation, RNAi was used to knock down the seventeen lipid genes in a *C. elegans* strain carrying a GFP reporter driven by three tandem repeats of the ESRE motif (*3XESRE*::GFP)^41^. This reporter was then exposed to rotenone, which is known to activate ESRE^42^. The majority of the knockdowns did not affect expression of the *3XESRE*::GFP reporter; *acdh-1(RNAi)* virtually abolished GFP expression in the presence of rotenone, while *emb-8(RNAi)* enhanced it **(Fig. 5A-B, Fig. S2A-D)**. Native ESRE gene expression was evaluated by qRT-PCR on a *C. elegans acs-5* mutant^43^, and no effect on expression was observed, validating our reporter results **(Fig. 5C)**.

**Figure 5.**
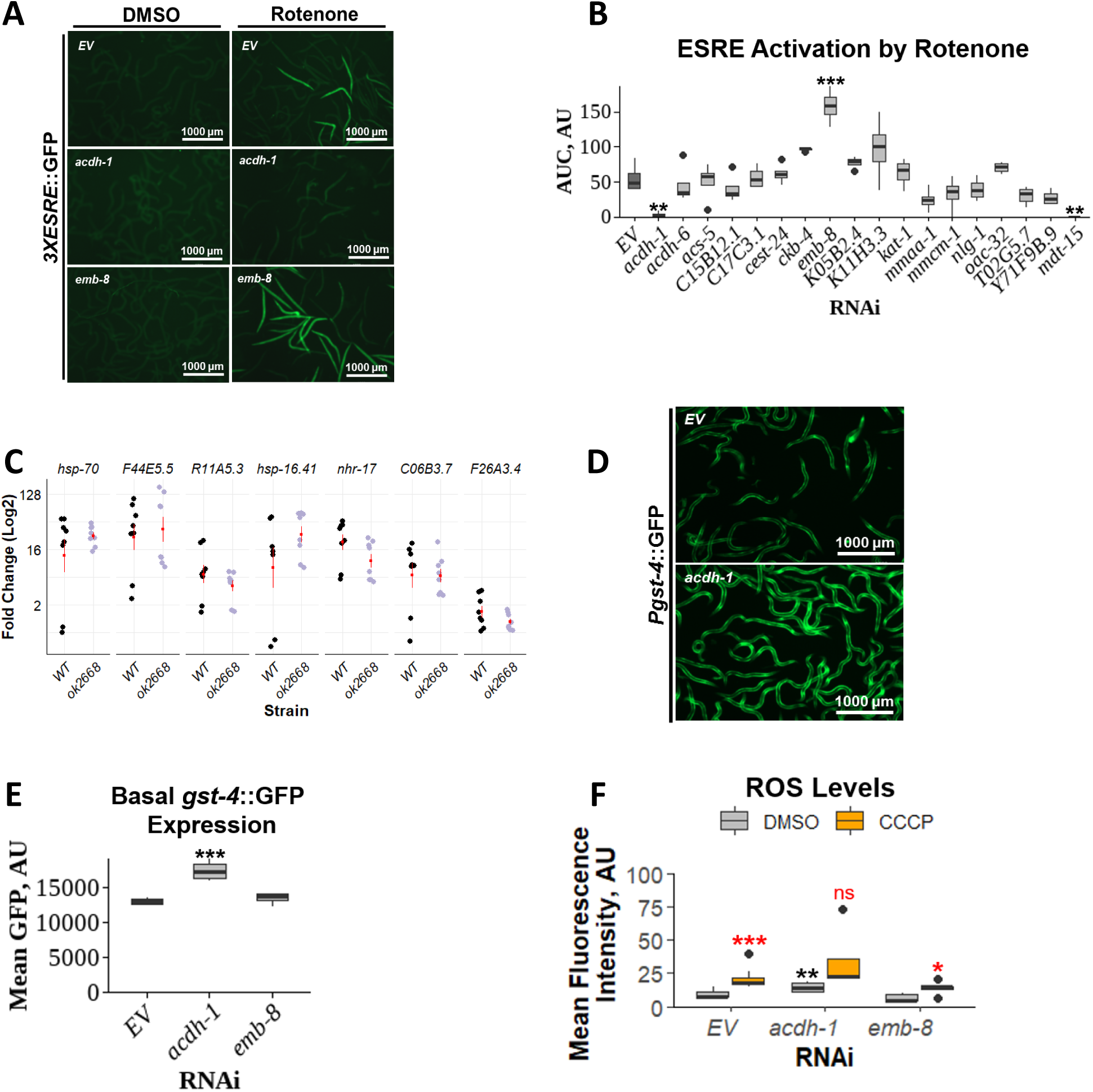
Lipid genes do not promote host defense via ESRE pathway activation. **(A)** Fluorescent images of *3XESRE*::GFP worms reared on *EV*, *acdh-1*, or *emb-8* RNAis and treated with 25 μM rotenone or *v/v* DMSO. **(B)** Area under the curve quantification for ESRE activation by rotenone across various hours. **(C)** Rotenone-induced native ESRE gene expression in *WT* or *acs-5(ok2668)* worms. Gene expression levels were normalized to each cohort’s DMSO control. **(D)** Fluorescent images of *gst-4p*::GFP worms reared on *EV* or *acdh-1* RNAis. **(E)** Quantification of mean GFP, which gives a readout on oxidative stress activation. **(F)** Quantification of basal and CCCP-induced ROS levels of *acdh-1* and *emb-8* knockdowns. Black stars indicate one-way ANOVA comparisons to the *EV(RNAi)* control within basal (DMSO) conditions. Red stars indicate paired comparisons between basal and induced ROS levels for each knockdown. A *t*-test was performed to calculate the significance of a treatment or condition when there were only two cohorts in the experimental setting. One-way analysis of variance (ANOVA) was performed to calculate the significance of a treatment or a condition when there were more than two cohorts in the experimental setting. No stars = not significant, * *p* < 0.05, ** *p* < 0.01, and *** *p* < 0.001.

To investigate if ACDH-1 and EMB-8 are involved in the direct signaling for ESRE activation, a *C. elegans* strain carrying a *gst-4*::GFP construct was used to evaluate their effect on oxidative stress^44^. Knockdown of *acdh-1* led to increased basal levels of *gst-4*::GFP, while *emb-8(RNAi)* showed no effect **(Fig. 5D-E)**. Levels of basal and CCCP-induced ROS were also quantified in these knockdowns, and the only observed effect was an increase in basal ROS levels by *acdh-1(RNAi)* **(Fig. 5F)**. Altogether, these data suggest that *acdh-1* and *emb-8* are more likely to be involved in the direct signaling required for ESRE activation, rather than indirect regulation by increasing ROS.

The role of these seventeen lipid genes on the activation of organellar proteostasis pathways (UPR^mt^) was evaluated by performing analogous experiments with an *hsp-6*::GFP reporter. *spg-7(RNAi)* was used to activate the UPR^mt^ pathway^45^ **(Fig. S3 A-B)**. None of the knockdowns affected UPR^mt^ activation. Given that LK-*Pa* triggers activation of chaperones to restore mitochondrial proteostasis, we evaluated its effect on endoplasmic reticulum (ER) proteostasis using an *hsp-4*::GFP reporter for UPR^ER^ **(Fig. S3C)**^46^. Our results indicate that *P. aeruginosa* triggers UPR^ER^ activation in a liquid and agar-based context **(Fig. S3D)**. To investigate whether lipid metabolism mediated the UPR^ER^ response to LK-*Pa*, *hsp-4*::GFP worms were fed RNAi targeting our seventeen lipid hits and then treated with tunicamycin, a *bona fide* UPR^ER^ activator^46^ **(Fig. S3E)**. Knockdown of *kat-1*, *oac-32*, and *Y71F9B.9* resulted in decreased UPR^ER^ activation **(Fig. S3F)**, while *mdt-15(RNAi)* increased UPR^ER^ activation, which is consistent with previous studies^19^.

Lastly, we studied the effect of our lipid gene hits on mitophagic and downstream autophagic activation by treatment with *trans*-β-nitrostyrene, a robust mitophagic activator in both human cells and *C. elegans*^47,48^. To test these, *C. elegans* strains carrying *pink-1*::PINK-1::GFP (mitophagy) or *lgg-1*::LGG-1::GFP (autophagy) translational fusions were used^48–50^. Out of the seventeen genes tested, five increased PINK-1::GFP levels **(Fig. 6A-B, Fig. S4 A-D)**. These five knockdowns were subsequently evaluated for their effect on LGG-1::GFP puncta levels **(Fig. 6C-D)**. Our results indicate that disrupting *acdh-6* or *C17C3.1* increased LGG-1::GFP puncta. This is in contrast to *emb-8(RNAi)*, which decreases the number of LGG-1::GFP puncta. Altogether, our data suggest a partial overlap of host resistance mediated by lipid metabolism genes with genes that regulate organellar surveillance.

**Figure 6.**
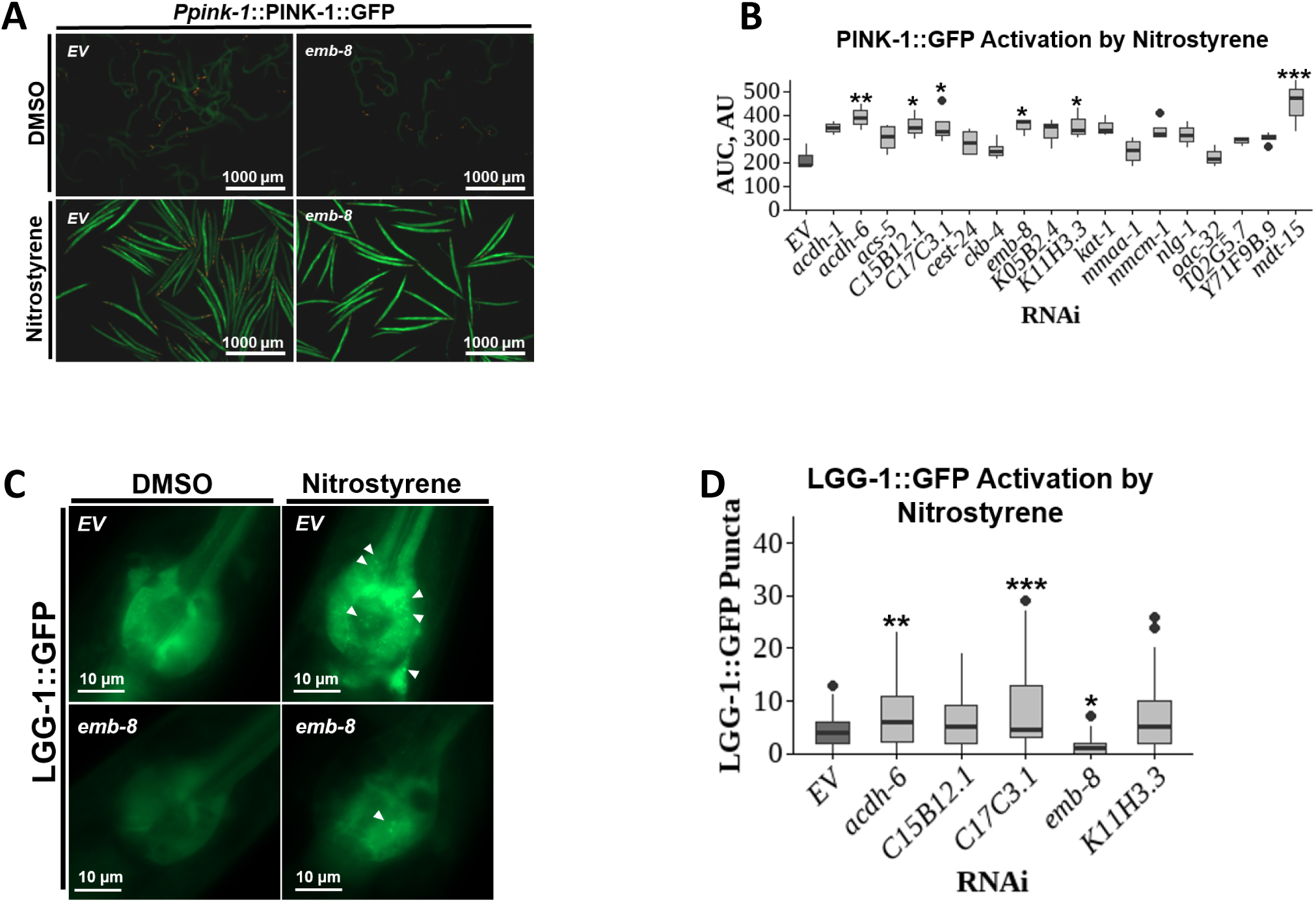
Lipid genes modulate mitophagic and downstream autophagic activation. **(A)** Fluorescent images of PINK-1::GFP worms reared on *EV* or *emb-8* RNAis and treated with 30 μM nitrostyrene or *v/v* DMSO. **(B)** Area under the curve quantification for PINK-1::GFP activation by nitrostyrene across various hours. **(C)** Fluorescent images of LGG-1::GFP worms reared on *EV* or *emb-8* RNAis and treated with 30 μM nitrostyrene or *v/v* DMSO. White arrows point to LGG-1 puncta in the worm’s pharyngeal terminal bulb. **(D)** Quantification of LGG-1 puncta. One-way analysis of variance (ANOVA) was performed to calculate the significance of a treatment or a condition when there were more than two cohorts in the experimental setting. No stars = not significant, * *p* < 0.05, ** *p* < 0.01, and *** *p* < 0.001.

### P. aeruginosa liquid-based pathogenesis triggers HLH-30-dependent metabolic rewiring

Our lab previously demonstrated that *P. aeruginosa* inefficiently colonizes *C. elegans* in a liquid setting^51^. This may be a consequence of diminished food uptake in liquid, which may limit the number of bacteria that enter the alimentary space. The transition to a liquid environment also reduces food uptake, which could alter the host’s metabolism as well^51^. Importantly, our lab showed that in the LK-*Pa* condition, *C. elegans* enters a dormant-like state, which is likely to serve a defensive function during exposure^51^.

Many animals activate and increase autophagic and lysosomal function during nutrient-scarce periods^52^. *C. elegans,* for example, activates the basic helix-loop-helix transcription factor HLH-30, which is out-competed by the Max-like factor MXL-3 when food is plentiful^53^. Since reduced metabolic activity and increased autophagy are associated with LK-*Pa,* we hypothesized that HLH-30 may be activated. To test this, a *C. elegans* strain carrying an *hlh-30*::HLH-30::GFP translational fusion was used^54^. Our results show that LK-*Pa* triggered an increase in nuclear localization of HLH-30 when compared to LK-*Ec* **(Fig. 7A-B, Fig. S6A)**. This suggested that HLH-30 activation may serve a protective role. We measured host survival of *hlh-30(tm1978)*^55^ worms in LK-*Pa* conditions and observed a significant increase in death **(Fig. 7C)**, confirming that HLH-30 plays a role in host defense in this context. HLH-30 loss-of-function did not affect worm survival in LK-*Ec* conditions **(Fig. S6B)**, proving that this outcome is due to the presence of the pathogen rather than non-specific sickness. Similar results were observed when exposing *hlh-30(RNAi)* worms in LK-*Pa* and LK-*Ec* conditions, validating these findings (Fig. S6C-D).

**Figure 7.**
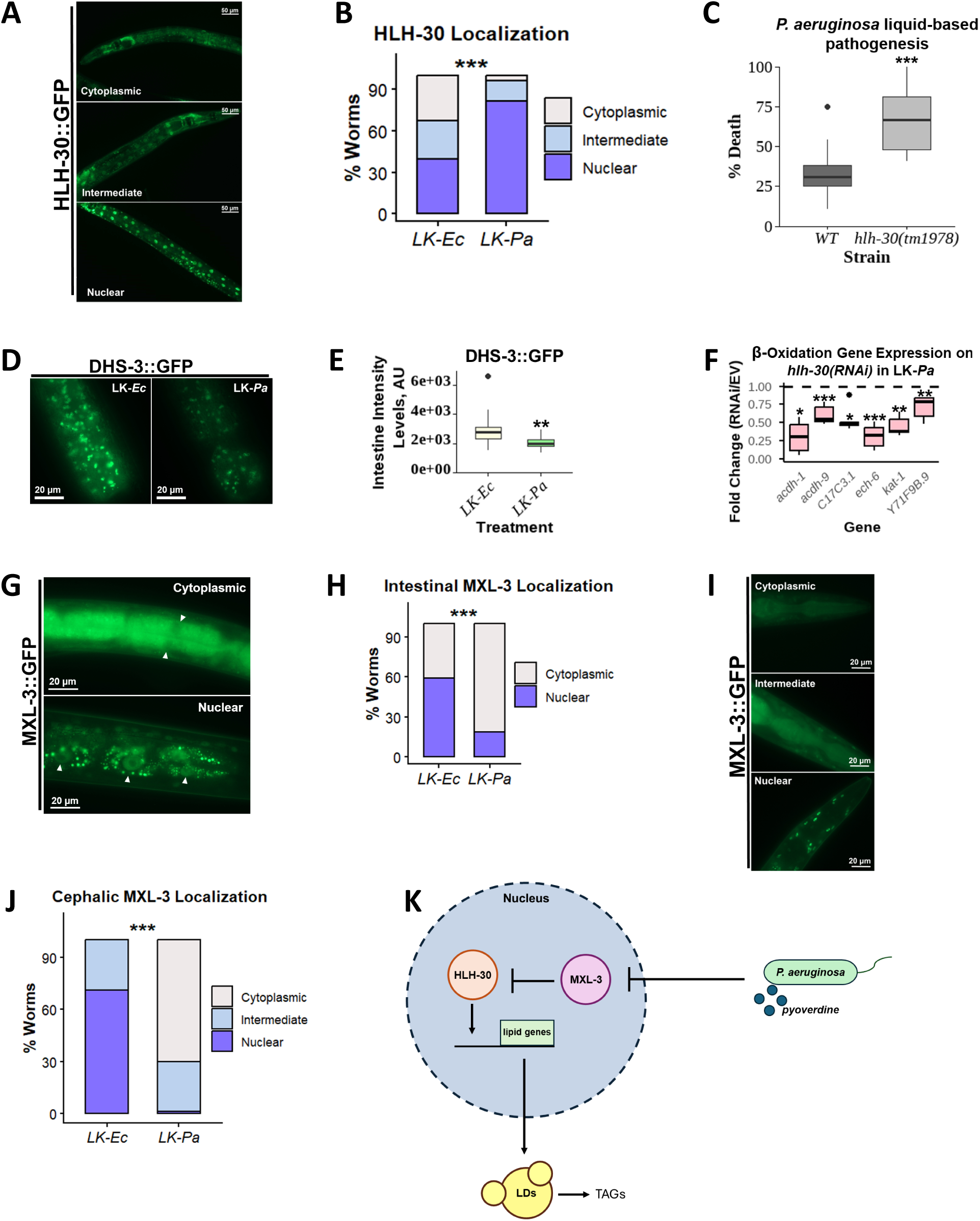
*P. aeruginosa* pathogenesis triggers HLH-30-dependent metabolic rewiring. **(A, G, I)** Representative images illustrating the scoring criteria used to classify HLH-30::GFP **(A)** or MXL-3::GFP **(G, I)** cellular localization as cytoplasmic, intermediate, or nuclear. **(B, H, J)** Quantification of HLH-30::GFP **(B)** or MXL-3::GFP **(H, J)** localization in worms exposed to LK-*Ec* or LK-*Pa*. Statistical significance was determined using a χ² test comparing localization distributions between treatments, * *p* < 0.05, ** *p* < 0.01, and *** *p* < 0.001. **(C)** Quantification of the percentage of dead WT or *hlh-30(1978)* worms exposed to LK-*Pa*. **(D)** Intestinal fluorescent images of DHS-3::GFP worms exposed to LK-*Ec* or LK-*Pa*. **(E)** Quantification of intestinal intensity DHS-3::GFP levels, which give a readout on intestinal lipid droplet content. **(F)** Native β-oxidation gene expression of worms fed with *EV(RNAi)* or *hlh-30(RNAi)* and exposed to LK-*Pa*. Gene expression levels were normalized to the *EV(RNAi)* control. A *t*-test was performed to calculate the significance of a treatment or condition when there were only two cohorts in the experimental setting. One-way analysis of variance (ANOVA) was performed to calculate the significance of a treatment or a condition when there were more than two cohorts in the experimental setting. No stars = not significant, * *p* < 0.05, ** *p* < 0.01, and *** *p* < 0.001.

Since LK-*Pa* triggered an increase in β-oxidation gene expression, we hypothesized that LK-*Pa* pathogenesis could also affect host lipid droplet content. To test this, a *C. elegans* strain carrying a *dhs-3*::DHS-3::GFP translational fusion was used^56^. We observed decreased intestinal DHS-3::GFP expression under LK-*Pa* conditions **(Fig. 7D-E)**. This both explains the enrichment of β-oxidation genes and highlights the downstream effects of HLH-30 activation. Next, we assessed the importance of host lipid droplet homeostasis in LK-*Pa* by testing survival of *lpin-1(RNAi)* worms. LPIN-1 is a key regulator for phosphatidic acid metabolism and lipid droplet/TAG formation, pathways whose function is vital for host lifespan^57^. Disruption of *lpin-1* increased host death during LK-*Pa* exposure with feeding on RNAi starting at the L3 stage; at the same time, survival in the liquid *E. coli* treatment did not result in significant changes in viability **(Fig. S6E-F).** This highlighted the importance of lipid homeostasis in the LK-*Pa* condition.

The effect of *hlh-30(RNAi)* on β-oxidation gene expression during LK-*Pa* was also evaluated. A significant decrease in expression of these genes was observed, indicating that HLH-30 activation drives their expression **(Fig. 7F)**. A comparison of previously published *hlh-30*-dependent genes and our seventeen lipid hits revealed that roughly 30% of them are known to be regulated by HLH-30^58^.

Because LK-*Pa* is a nutrient-poor condition, we hypothesized that HLH-30 activation could be due to it outcompeting MXL-3 at the promoters of target genes. To test this, a *C. elegans* strain carrying an MXL-3::TY1::EGFP::3xFLAG translational fusion was used^59,60^. Our results show that in contrast to the HLH-30::GFP response to LK-*Pa*, intestinal and cephalic MXL-3 remained cytoplasmic, while in LK-*Ec*, it translocated into the nucleus **(Fig. 7G-J, Fig. S6G-H)**. Altogether, these results uncover a regulatory mechanism in which LK-*Pa* condition prevent nuclear localization of MXL-3, permitting HLH-30 activation and downstream lipid droplet degradation and β-oxidation **(Fig. 7K)**.

## Discussion

In this study, we demonstrate that lipid metabolism is a major component of the host defense response against liquid-based *P. aeruginosa* pathogenesis. Our data show that the host metabolic response to liquid-based infection is largely driven by pyoverdine-mediated iron deprivation and differs from that of agar-based infection. Through RNAi screening, we identified seventeen lipid genes crucial for mounting the host defense response against liquid-based infection by Gram-negative and Gram-positive pathogens.

Previous research by O’Rourke and Ruvkun demonstrated that HLH-30 and MXL-3 function as opposing regulators for the mobilization of lipids in low-nutrient environments^53^. Upon localizing to the nucleus, MXL-3 downregulates lysosomal lipases, whereas cytoplasmic localization allows HLH-30 to accumulate in the nucleus and promote expression of lysosomal lipases and autophagy genes, enabling the utilization of cellular lipid stores^53^. Our findings extend this nutrient-responsive pathway to host defense against *P. aeruginosa* acute infection. Using fluorescence-based reporters, we show that this metabolic response involved inhibition of MXL-3 nuclear translocation while activation of HLH-30 nuclear translocation, resulting in lipid droplet depletion, and increased β-oxidation. Notably, Goswamy et al. previously established HLH-30-mediated lipid droplet depletion as a host defense mechanism against *S. aureus* and *E. faecalis* infection^61^. Altogether, these studies, along with our findings, establish a relationship between pathogen-induced metabolic stress and innate immunity and reveal HLH-30-dependent metabolic rewiring as a critical strategy through which *C. elegans* adapts to LK-*Pa*.

In this study, we demonstrate that the metabolic response to acute infection functions largely independently of organellar surveillance pathways. However, our results suggest that pathogen-induced metabolic stress may be connected to endoplasmic reticulum (ER) homeostasis. Previous studies have shown that bacterial siderophores can disrupt host iron homeostasis and mitochondrial iron availability^30,39,62^. In LK-*Pa*, where pyoverdine-mediated iron sequestration impairs mitochondrial function and is associated with extensive metabolic remodeling, these perturbations in iron and cellular metabolism could consequently place additional demands on the ER. Consistent with a potential connection between lipid metabolism and ER homeostasis, our results showed that KAT-1, OAC-32, and Y71F9B.9 were required for UPR^ER^ activation following tunicamycin treatment. Our findings suggest that specific components of lipid metabolism may influence the capacity of the ER stress response. Whether lipid metabolic remodeling directly regulates ER stress during infection or instead represents a parallel response to a common upstream signal such as iron deprivation remains to be determined.

Notably, ultrastructural analyses via electron microscopy of animal models have shown that *E. faecalis* in sepsis damages host mitochondria^63^. Additionally, research using human cell lines has demonstrated that *S. aureus* releases extracellular vesicles that localize to host mitochondria, trigger mitochondrial damage, and prevent the recycling of damaged mitochondria via the PINK-1/Parkin pathway due to the acidic environment of these vesicles^64^. Since liquid-based infection by *E. faecalis* and *S. aureus* caused higher death in a large portion of lipid metabolism knockdowns we identified as playing a role in LK-*Pa*, we hypothesize these pathogens might trigger a similar metabolic rewiring. Research in our lab has established that liquid-based pathogenesis by *E. faecalis* or fungal pathogen *Candida albicans* exhibited significantly diminished host colonization^51^, suggesting similar trends for liquid-based pathogenesis models. It is also critical to evaluate the localization profiles of HLH-30 and MXL-3 in these pathosystems, as liquid-based models reduce the uptake of nutrients. However, most crucial of all is performing transcriptional profiling of *C. elegans* exposed to LK-*Ef* and LK-*Sa*. Doing so will reveal the central components involved in promoting host adaptation against liquid-based pathogenesis by *E. faecalis* and *S. aureus*.

## Materials and Methods

### C. elegans strains

All *C. elegans* strains were maintained on nematode growth medium (NGM) plates seeded with *E. coli* OP50 as the food source and were maintained at 22°C^65^, unless noted otherwise. *C. elegans* strains used include: N2 Bristol (wild-type)^66^, SS104 |*glp-4(bn2)I*|^67^, WY703 |*fdIs2*[*3XESRE*::GFP]; pFF4[*rol-6(su1006)*]|^68^, CL2166 |*dvIs19*[(pAF15)*gst-4p*::GFP::NLS]|^44^, RB2015 |*acs-5(ok2668)* III|^43^, SJ4100 |*zcIs13*[*Phsp-6*::GFP]|^45^, SJ4005 |zcIs4 [*hsp-4p*::GFP] V|^69^, NVK90 |*pink-1*(*tm1779*); *houIs001*{*byEx655* [*pink-1p*::PINK-1::GFP + *myo*-2*p*::mCherry]}^48^, DA2123 |*adIs2122*[*lgg-1p*::GFP::LGG-1 + *rol-6*(*su1006*)]|^49,50^, MAH240 |sqIs17 [*hlh-30p*::HLH-30::GFP + *rol-6*(*su1006*)]|^54^, JIN1375 |*hlh-30(tm1978)* IV|^55^, LIU1 |drIs1 [*dhs-3p*::DHS-3::GFP + *unc-76*(+)]|^56^, OP745 |wgIs745 [*mxl-3p*::MXL-3::TY1::EGFP::3xFLAG + *unc-119*(+)]|^59,60^.

Synchronized worms were prepared by hypochlorite isolation of eggs from gravid adults, followed by hatching of the embryos in pure S Basal. 6,000 synchronized L1 larvae were reared on 10 cm NGM plates seeded with OP50. After transfer, worms were grown at 22°C for 48 hours before experiments, or three days for the next isolation of eggs. For synchronization of SS104 worms, 6,000 synchronized L1 larvae were reared on 10 cm NGM plates seeded with OP50 and grown at 15°C for five days for the next isolation of eggs. L4-stage hermaphrodite worms were used for all assays unless specified in the text.

### Bacterial strains

RNAi experiments in this study were done using RNAi-competent *E. coli* (HT115-based) obtained from the Ahringer or Vidal RNAi libraries^70,71^. For experiments requiring OP50-based RNAi, plasmids isolated from RNAi-competent *E. coli* (HT115-based) were transformed into RNAi-competent *E. coli* OP50. All constructs were sequence-verified before use.

For pathogenesis assays, *P. aeruginosa* PA14^72,73^, methicillin-resistant *S. aureus* MRSA131^74^, and *E. faecalis* OG1RF^75^ were used. *P. aeruginosa* was cultured in Luria-Bertani (LB) medium, *S. aureus* in tryptic soy broth (TSB), and *E. faecalis* in brain heart infusion broth (BHI).

### RNA interference protocol

RNAi-expressing *E. coli* were cultured and seeded onto NGM plates supplemented with 25 μg/mL carbenicillin and 1 mM IPTG. For RNAi experiments starting at L1 or L3, 1,500 synchronized L1 or L3 larvae were reared on 6 cm RNAi plates and grown at 22°C for 48 hours before imaging, exposure to chemical treatment, or pathogens. For double RNAi experiments, specifically for *Phsp-6*::GFP experiments, RNAi of interest was mixed at a 3:1 ratio with *spg-7(RNAi)* or *EV(RNAi)* control, respectively.

### *P. aeruginosa* pathogenesis assays

*P. aeruginosa* liquid-based pathogenesis model (Liquid Killing or LK-*Pa*) was performed essentially as described^76,77^. 25 synchronized young adult *glp-4* mutants were sorted into each well of a 384-well plate. LK medium was mixed with *P. aeruginosa* PA14 and then added into each well (final OD_600_: 0.03). Plates were incubated at 25 °C. At different time points, plates were washed three times, and worms were stained with SYTOX Orange nucleic acid stain (Invitrogen) for 12 hours to identify dead worms. Afterwards, plates were washed and then imaged using a Cytation5 automated microscope, and the fraction of dead worms was quantified autonomously with Cell Profiler software^78^.

*P. aeruginosa* agar-based pathogenesis model (Slow Killing or SK-*Pa*) was performed as previously described^79,80^. 50 young adult worms were transferred onto SK-*Pa* plates and incubated at 25 °C. Worms were scored daily to obtain survival curves; dead worms were extracted from assay plates, and worms that left the surface of the plate were censored from data analysis.

*P. aeruginosa* liquid-based exposure of GFP-carrying worms was performed similarly to the LK-*Pa* assay. About 150 worms were sorted into each well of a 96-well full-area plate. Liquid Killing medium was mixed with *P. aeruginosa* PA14 and then added into each well for a total volume of 160 μL (final OD_600_: 0.03). After 24 hours, bacteria were washed off the plates, and worms were imaged.

### *P. aeruginosa*-derived pyoverdine-rich filtrate assay

*P. aeruginosa*-derived pyoverdine-rich filtrate was prepared as previously described^39,81,82^. In short, PA14 was first cultured in 5 mL of LB at 37°C with constant agitation. Then, saturated overnight culture was sub cultured into low-iron M9 media [(1% w/v) Difco 5X M9 salts, 11.3 g/L low-iron casamino acids, 0.4% glucose, 1mM CaCl_2_, 1mM Mg_2_SO_4_)] for 24h at 37°C with constant agitation. Bacteria were removed by centrifugation and subsequent filtration through a 0.2-mm filter. Pyoverdine concentration in the filtered sample was then measured fluorometrically, and only samples with high levels of pyoverdine were used in subsequent treatments. 25 synchronized young adult *glp-4* mutants were sorted into each well of a 384-well plate. Pyoverdine-rich filtrate was mixed with *E. coli* OP50 and then added into each well (final OD_600_: 0.03). Plates were incubated at 25 C° . At different time points, plates were washed three times with S Basal, and then worms were stained with SYTOX Orange nucleic acid stain (Invitrogen) for 12 hours to identify dead worms. Afterwards, plates were washed and then imaged using a Cytation5 automated microscope, and the fraction of dead worms was quantified autonomously with Cell Profiler software^78^.

### Enterococcus faecalis pathogenesis assays

*E. faecalis* liquid-based pathogenesis model (Liquid Killing or LK-*Ef*) was performed essentially as described^83^. 25 synchronized young adult *glp-4* mutants were sorted into each well of a 384-well plate. LK medium was mixed with *E. faecalis* OG1RF and then added into each well (10% BHI and final OD_600_: 0.03). Plates were incubated at 25 °C. At different time points, worms were transferred to new 384-well plates with S basal with 0.1% tween and then washed with S basal. Then, worms were stained with SYTOX Orange nucleic acid stain (Invitrogen) for 12 hours to identify dead worms. Afterwards, plates were washed and then imaged using a Cytation5 automated microscope, and the fraction of dead worms was quantified autonomously with Cell Profiler software^78^.

### Staphylococcus aureus pathogenesis assays

*S. aureus* liquid-based pathogenesis model (Liquid Killing or LK-*Sa*) was performed essentially as described^84^. 25 synchronized young adult *glp-4* mutants were sorted into each well of a 384-well plate. LK medium was mixed with *S. aureus* MRSA131 and then added into each well (10% TSB and final OD600: 0.03). Plates were incubated at 25 °C. At different time points, worms were transferred to new 384-well plates with S Basal with 0.1% Tween and then washed with S Basal. Then, worms were stained with SYTOX Orange nucleic acid stain (Invitrogen) for 12 hours to identify dead worms. Afterwards, plates were washed and then imaged using a Cytation5 automated microscope, and the fraction of dead worms was quantified autonomously with Cell Profiler software^78^.

### *C. elegans* chemical exposure assays

Synchronized L4-stage worms were washed from NGM plates seeded with *E. coli* OP50, HT115-based RNAi-competent *E. coli*, or OP50-based RNAi-competent *E. coli* into a 15 mL conical tube and rinsed three times. For experiments involving the *3XESRE*::GFP strain, worms were sorted into a 96-well half-area plate (∼100 worms/well), and the total volume per well, including treatment, was 100 µL. Treatment with rotenone was done at a final concentration of 25 µM per well. Treatment with nitrostyrene, was done at a final concentration of 30 µM per well. Treatment with tunicamycin was done at a final concentration of 60 µM per well. DMSO was used as the solvent control for all of these treatments at a *v/v* ratio relative to the treatment.

For experiments involving the use of the *gst-4p*::GFP, *hsp-4p*::GFP, PINK-1::GFP strains, worms were sorted into a 96-well full area plate (∼150 worms/well), and the total volume per well, including treatment, was 160 µL. Worms were imaged with a Cytation5 automated microscope every hour for eighteen hours at room temperature.

### ROS measurements

*C. elegans* ROS measurements were conducted as previously described, with some adjustments^85^. Synchronized L4-stage *glp-4* worms were washed from 6 cm NGM plates seeded with OP50-based RNAi-expressing *E. coli* into a 1.5 mL tube and rinsed three times with S Basal and 0.1% Tween. Worms were sorted into a 96-well half-area plate (∼50 worms/well), and the total volume per well, including treatment, was 100 µL.

To induce ROS, worms were treated with 78.2 µM CCCP, and to quantify/visualize ROS, worms were stained with a final concentration of 25 µM DCFDA per well. Worms were imaged, and fluorescence intensity was measured using a Cytation5 automated microscope at 0 hours, 4 hours, and 5 hours. Mean fluorescence intensity (MFI) was calculated by dividing the fluorescence intensity by the number of worms per well.

### High-throughput fluorescence imaging and quantification

Worms were imaged with a Cytation5 automated microscope every hour for eighteen hours at room temperature. Percent GFP area quantification was calculated by dividing the object sum area corresponding to “highly activated” GFP of worms in a well by the object sum area of all the worms within the well using Gen5 software. This was performed by setting up two “Cellular Analysis” calculations in Gen5; one calculates the object sum area of all the worms in a well by setting a low GFP channel threshold (highlights worms by their background fluorescence), and the other one calculates the object sum area of the “highly activated” GFP portions of worms in the well by setting a high GFP channel threshold.

### Quantitative reverse transcriptase PCR (qRT-PCR)

30,000 worms were used for RNA extraction, and subsequent qRT-PCR was performed as previously described^76^. Before RNA extraction, worms were reared on RNAi plates starting at the L1 stage. For treatment with rotenone, L4-stage worms were washed off plates, rinsed three times, and incubated for 8 hours in S Basal supplemented with 25 μM rotenone or a corresponding volume of DMSO.

For LK-*Pa* or LK-*Ec* treatment, young adult *glp-4* worms were washed off plates, rinsed three times, and sorted onto a 6-well plate (∼10,000 worms/mL per well). Then, 2 mL of LK medium containing either *P. aeruginosa* PA14 or *E. coli* OP50 was added to each well, yielding a final volume of 3 mL and a final OD_600_ of 0.03. Worms were incubated in the treatment for 24 hours at 25°C before RNA extraction.

To generate cDNA, a total of 1.5 μg of total RNA was used (NEB LunaScript RT SuperMix Kit). qRT-PCR was performed with NEB Luna® Universal qPCR Master Mix using a Bio-Rad CFX96 system. Fold changes were derived using the ΔΔC_t_ method. At least three biological replicates were used for statistical analyses.

### Microscopy

For visualization of DA2123 (LGG-1::GFP), OP745 (MXL-3::GFP), and LIU1 (DHS-3::GFP), worms were immobilized using 10 mM levamisole and transferred onto 3% noble agar slides. At least three biological replicates with 15 worms per replicate were imaged using a Zeiss ApoTome.2 Imager.M2, Carl Zeiss, Germany with 63x magnification. For OP745, nuclei in the head were imaged. For LIU1, lipid droplets in the intestine were imaged. For visualization of MAH240, worms were imaged using a Zeiss ApoTome.2 Imager.M2, Carl Zeiss, Germany with 20x magnification.

Intestinal fluorescence intensity of LIU1 was quantified using the surfaces tool on a 10.2.0 version of Imaris software. Parameters were stored for batch to ensure each image was analyzed consistently. For DA2123, fluorescent puncta in the terminal bulb of the pharynx were imaged. The terminal pharyngeal bulb was set as the ROI, and LGG-1 puncta were counted using the spots tool on a 10.2.0 version of Imaris software.

## Statistical analyses

At least three biological replicates were performed for each experiment. RStudio (version 4.3.0) was used to perform all statistical analyses. A *t*-test was performed to calculate the significance of a treatment or condition when there were only two cohorts in the experimental setting. One-way analysis of variance (ANOVA) was performed to calculate the significance of a treatment or a condition when there were three or more cohorts in the experimental setting. Statistically significant results, as determined via ANOVA, were then followed by the *post-hoc* Dunnett’s test to calculate statistical significance or *p* values between each group compared to the control group. Statistical significance is shown in graphs as follows: no stars = not significant, \**p* < 0.05, ** *p* < 0.01, and *** *p* < 0.001.

Pearson’s chi-square test of independence was performed to assess the differences in categorical data. To identify which categories contributed the most to the overall association, standardized Pearson residuals were calculated for each cell in the contingency table. Cell-specific two-sided *p*-values were derived from the standardized residuals and were adjusted for multiple comparisons using the Benjamini-Hochberg false discovery rate (FDR) correction. Heatmaps were generated using standardized residuals, where positive residuals indicate over-representation and negative residuals indicate under-representation relative to expected counts under the null hypothesis of independence.

Spearman’s rank correlation analysis was used to assess the relationship between susceptibility phenotypes across different pathogenesis models. Statistical significance was calculated by using a two-sided Spearman correlation test. The resulting *p*-value indicates whether the observed correlation between two pathogenesis models is likely due to shared susceptibility patterns rather than pure chance.

## Supporting information

Combined Supplemental Figures and Tables

## Acknowledgements

Some strains were provided by the CGC, which is funded by NIH Office of Research Infrastructure Programs (P40 OD010440). This research was supported by NIH NIGMS R35GM129294 grant to NVK and a supplement for NVK’s NCI award R21CA280500 to LA (3R21CA280500-01A1S1) and training grant T32 AI055449-17 to LA. FV was supported by the Fulbright fellowship and YA was supported by NSF REU award 2244041.

## Author contributions

Conceptualization, NVK, LA, ET; data acquisition, LA, ET, YA, FV, AS, JL, NS, AVR, AH, EP; formal analysis, NVK, LA, ET, YA, FV; writing – original draft, NVK, LA, ET; writing – review and editing, all authors; visualization, LA, ET, YA, FV; supervision, NVK; funding acquisition, NVK.

## Supplementary Figure Legends

**Figure S1. Non-pathogenic conditions induce host baseline metabolism changes**

**(A-B)** Wormcat gene enrichment analyses of genes upregulated in response to LK-*Ec* **(A)** or SK-*Ec* **(B)**.

**Figure S2. Knockdown of *acdh-1* or *emb-8* affects ESRE activation**

**(A-D)** Curves representing ESRE activation by rotenone across over time.

**Figure S3. *P. aeruginosa* infection triggers UPR^ER^ activation, and lipid genes affect ER but not mitochondrial proteostasis**

**(A)** Fluorescent images of *hsp-6p*::GFP worms reared on *EV*, *atfs-1*, or *acdh-1* RNAis. Each RNAi was mixed in a 3:1 ratio with *spg-7* or *EV* RNAis. **(B)** Quantification of percent GFP area for specified condition, which gives a readout on UPR^mt^ activation. **(C)** Fluorescent images of *hsp-4p*::GFP worms exposed to LK-*Ec*, LK-*Pa*, SK-*Ec*, or SK-*Pa*. **(D)** Quantification of mean GFP, which gives a readout on UPR^ER^ activation. **(E)** Fluorescent images of *hsp-4p*::GFP worms reared on *EV*, *mdt-15*, *kat-1*, or *oac-32* RNAis and treated with 60 μM tunicamycin. **(F)** Quantification of mean GFP, which gives a readout on UPR^ER^ activation by tunicamycin. A *t*-test was performed to calculate the significance of a treatment or condition when there were only two cohorts in the experimental setting. One-way analysis of variance (ANOVA) was performed to calculate the significance of a treatment or a condition when there were more than two cohorts in the experimental setting. No stars = not significant, * *p* < 0.05, ** *p* < 0.01, and *** *p* < 0.001.

**Figure S4. Knockdown of *mdt-15*, *kat-1*, *Y71F9B.9*, or *oac-32* affects UPR^ER^ activation upon tunicamycin treatment**

**(A-D)** Curves representing UPR^ER^ activation by tunicamycin across various hours.

**Figure S5. Knockdown of *acdh-6*, *C15B12.1*, *C17C3.1*, *emb-8*, or *K11H3.3* affects PINK-1::GFP activation**

**(A-D)** Curves representing PINK-1::GFP activation by nitrostyrene over time.

**Figure S6. *P. aeruginosa* liquid-based pathogenesis triggers HLH-30-dependent metabolic rewiring**

**(A, G, H)** Standardized residual heatmaps from the χ² analyses on HLH-30::GFP **(A)** or intestinal **(G)** and cephalic MXL-3::GFP **(H)**. Positive residuals (red) indicate categories occurring more frequently than expected. Negative residuals (blue) indicate categories occurring less frequently than expected, \**p* < 0.05, ** *p* < 0.01, and *** *p* < 0.001. **(B-F)** Quantification of the percentage of dead worms upon treatment with specified condition. A *t*-test was performed to calculate the significance of a treatment or condition when there were only two cohorts in the experimental setting. No stars = not significant and * *p* < 0.05.

## Supplementary Tables

