## Supplementary material for "*P. aeruginosa* liquid-based pathogenesis triggers HLH-30-dependent metabolic rewiring in *C. elegans*": Combined Supplemental Figures and Tables

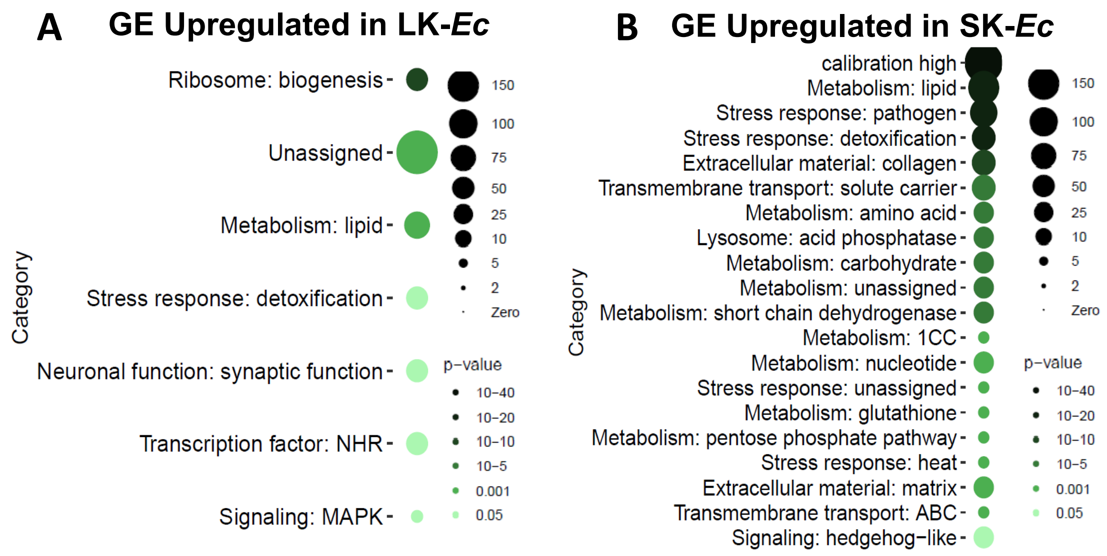


Supplementary Figure S1

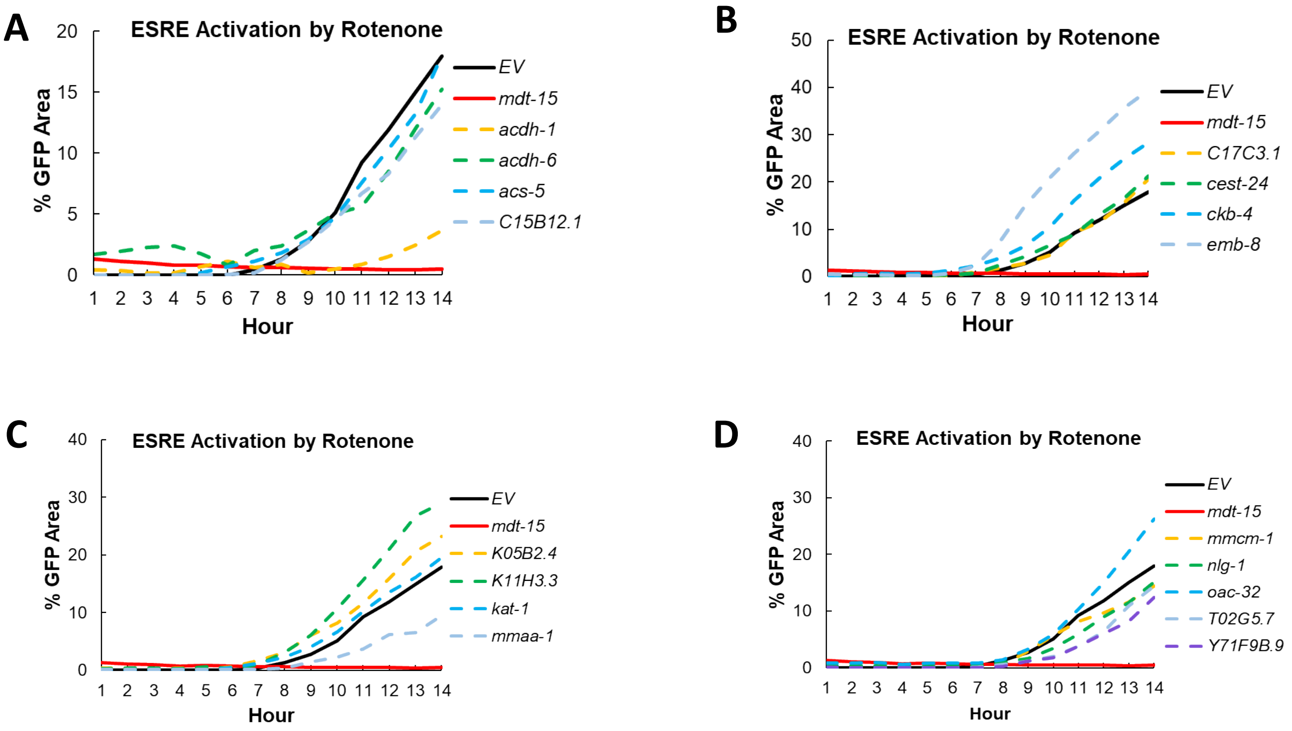


Supplementary Figure S2

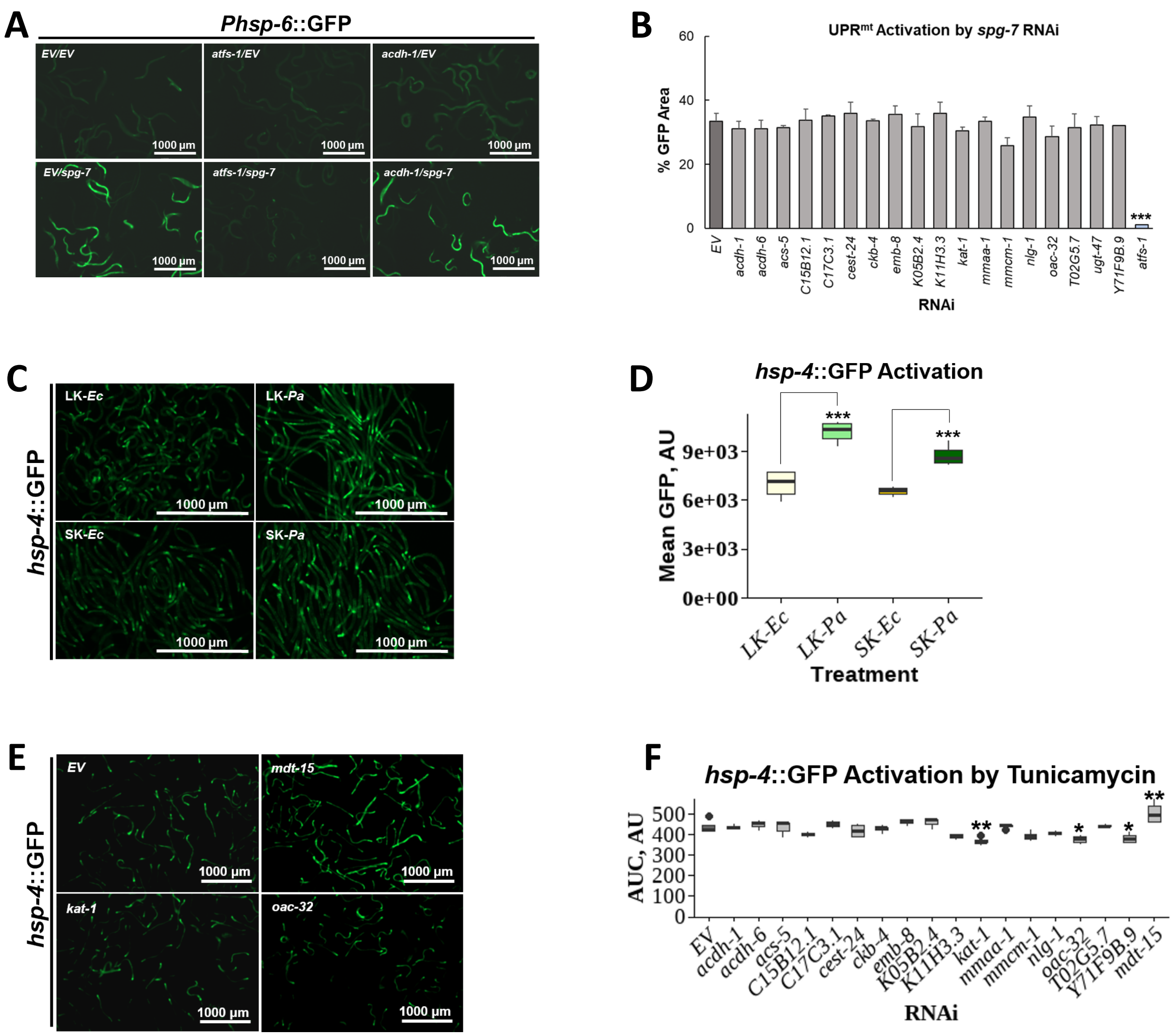


Supplementary Figure S3

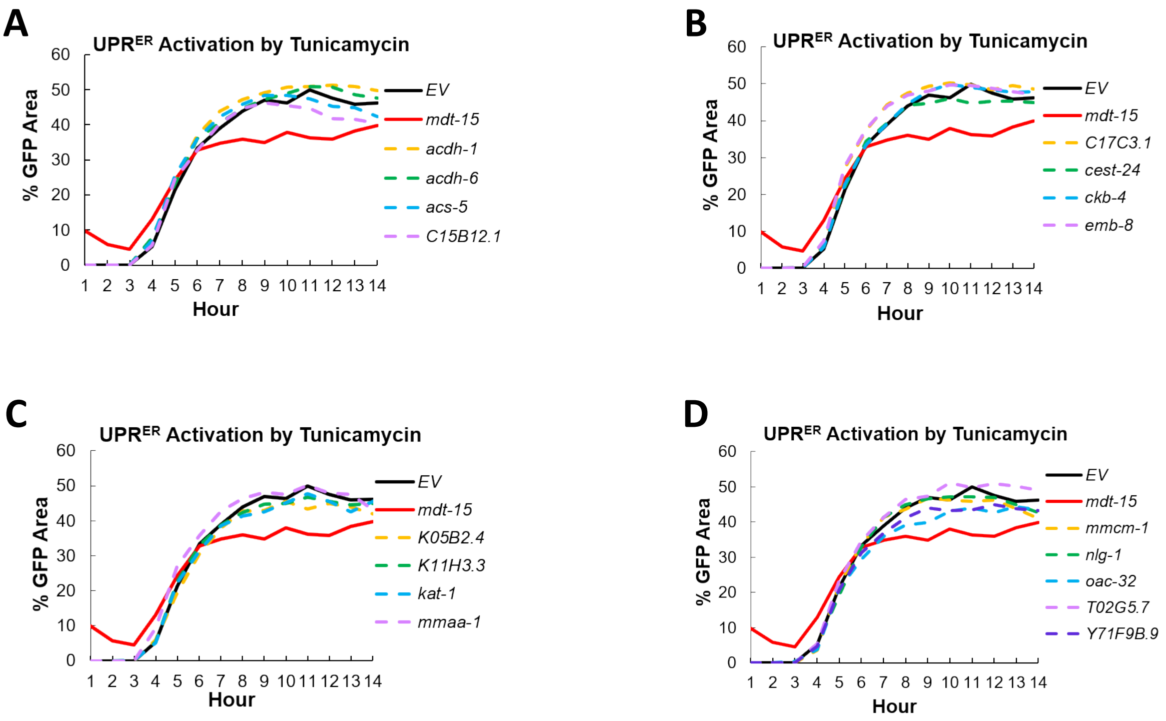


Supplementary Figure S4

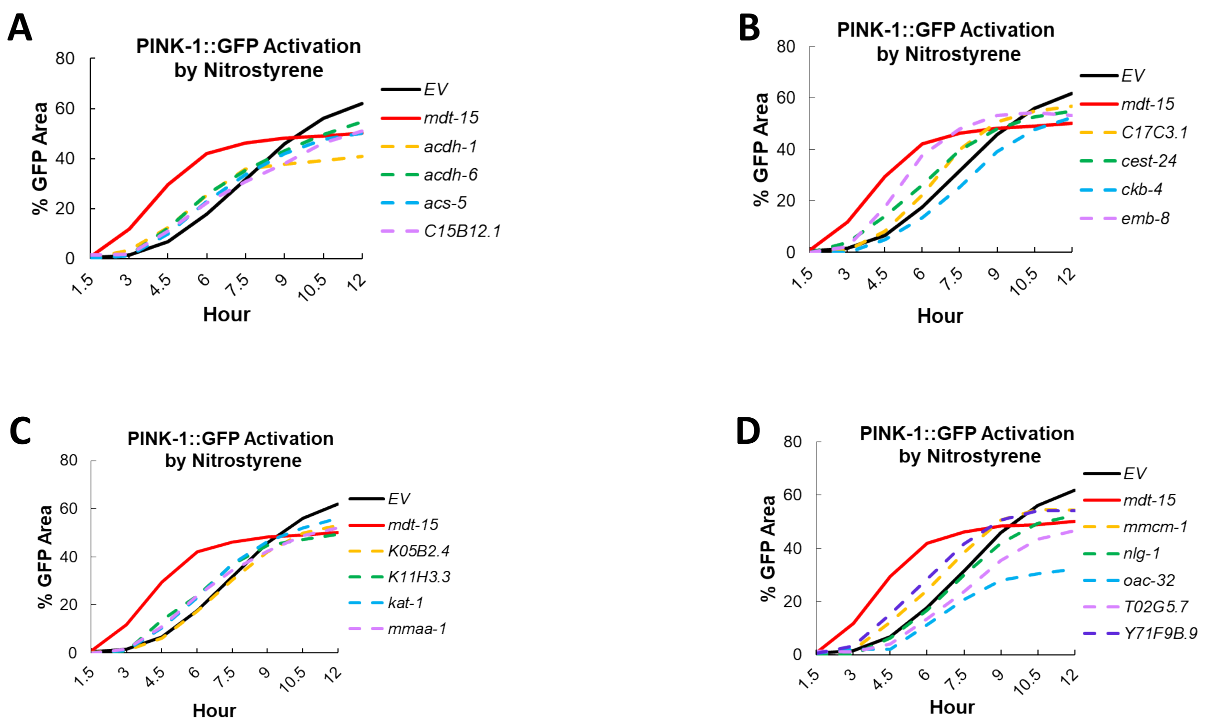


Supplementary Figure S5

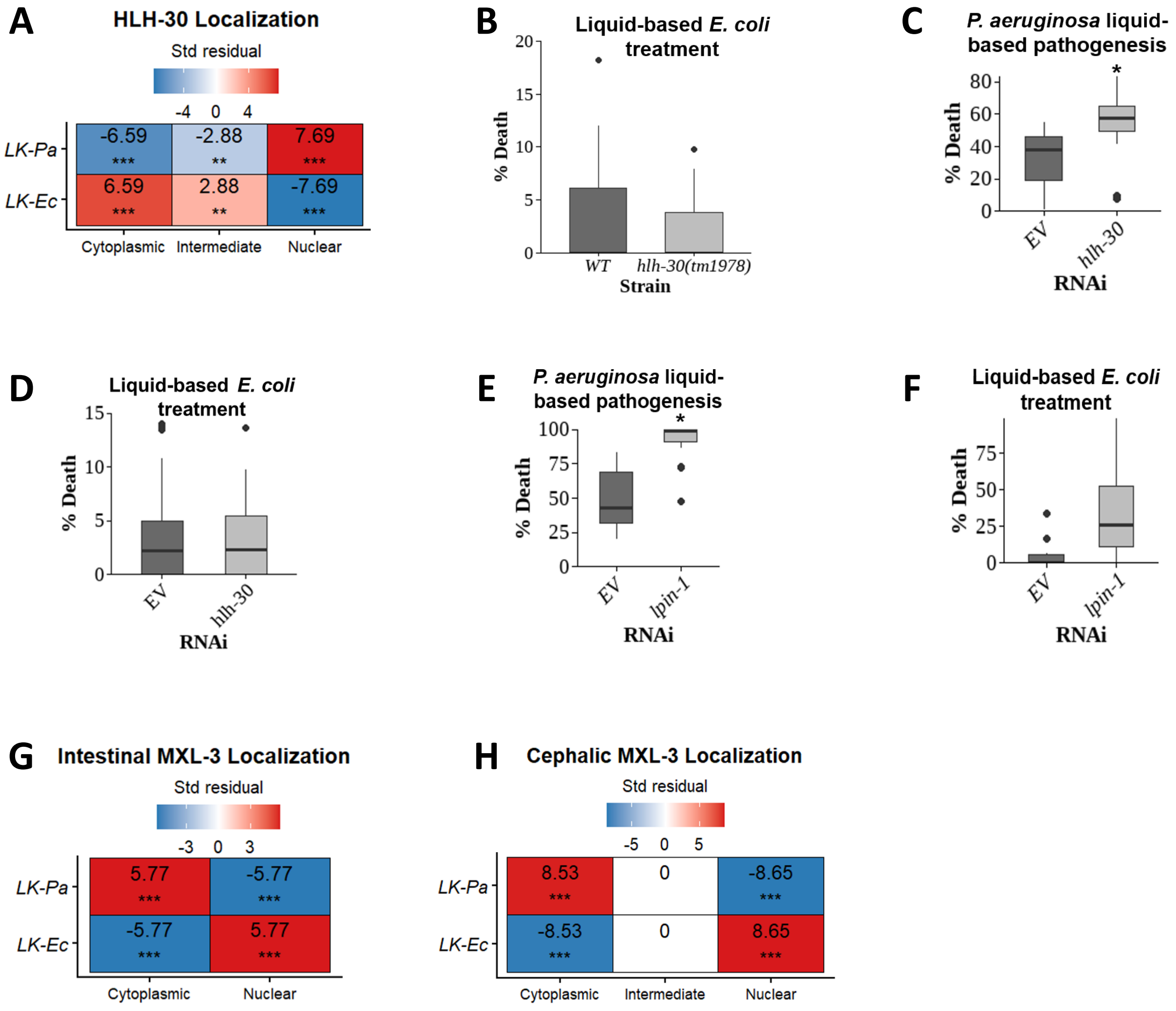


Supplementary Figure S6

Supplementary Table S1.

**Enriched GE category for top 300 genes in PC1 (negative)**

| Category | p-value | Bonferroni |
|---|---|---|
| Extracellular material: collagen | 1.51E-09 | 1.51E-07 |
| Metabolism: amino acid: synthesis | 2.81E-07 | 2.81E-05 |
| Metabolism: carbohydrate | 2.94E-06 | 0.000294 |
| Metabolism: pentose phosphate pathway | 1.76E-05 | 0.001764 |
| Metabolism: lipid: beta oxidation | 1.86E-05 | 0.001863 |
| Stress response: detoxification: GST | 1.99E-05 | 0.001991 |
| Metabolism: nucleotide | 2.13E-05 | 0.002133 |
| Lysosome: acid phosphatase | 2.31E-05 | 0.00231 |
| Metabolism: carbonic anhydrase | 6.89E-05 | 0.006894 |
| Proteolysis general: unassigned | 7.99E-05 | 0.007993 |

**Enriched GE category for top 300 genes in PC1 (positive)**

| Category | p-value | Bonferroni |
|---|---|---|
| Ribosome: biogenesis | 1.43E-20 | 1.58E-18 |
| Non-coding RNA: tRNA: production | 7.85E-05 | 0.008637 |

**Enriched GE category for top 300 genes in PC2 (negative)**

| Category | p-value | Bonferroni |
|---|---|---|
| Metabolism: lipid: beta oxidation | 1.42E-08 | 1.69E-06 |
| Metabolism: amino acid: breakdown | 4.28E-05 | 0.005091 |
| Metabolism: FMO | 4.54E-05 | 0.005402 |

**Enriched GE category for top 300 genes in PC2 (positive)**

| Category | p-value | Bonferroni |
|---|---|---|
| Stress response: pathogen: unassigned | 6.12E-09 | 6.49E-07 |
| Protein modification: carbohydrate | 6.56E-06 | 0.000695 |
| Unassigned: regulated by multiple stresses | 1.87E-05 | 0.001983 |
| Transmembrane transport: major facilitator | 3.33E-05 | 0.003531 |
